# TET proteins regulate Drosha expression and impact microRNAs in iNKT cells

**DOI:** 10.1101/2024.07.31.605991

**Authors:** Marianthi Gioulbasani, Tarmo Äijö, Jair E. Valenzuela, Julia Buquera Bettes, Ageliki Tsagaratou

**Affiliations:** Lineberger Comprehensive Cancer Center, University of North Carolina at Chapel Hill, Chapel Hill, NC, USA; School of Biology, Aristotle University of Thessaloniki, 54124 Thessaloniki, Greece; Joint Department of Biomedical Engineering, University of North Carolina at Chapel Hill and North Carolina State University, Raleigh, NC, USA; Department of Genetics, University of North Carolina at Chapel Hill, Chapel Hill, NC, USA; Department of Microbiology and Immunology, University of North Carolina at Chapel Hill, Chapel Hill, NC, USA

**Author notes:** These authors contributed equally to this work and share first co-authorship (Marianthi Gioulbasani, Tarmo Äijö).

**Keywords:** TET proteins, 5hmC, iNKT, Drosha, microRNAs, PLZF

## Abstract

DNA demethylases TET2 and TET3 play a fundamental role in thymic invariant natural killer T (iNKT) cell differentiation by mediating DNA demethylation of genes encoding for lineage specifying factors. Paradoxically, differential gene expression analysis revealed that significant number of genes were upregulated upon TET2 and TET3 loss in iNKT cells. This unexpected finding could be potentially explained if loss of TET proteins was reducing the expression of proteins that suppress gene expression. In this study, we discover that TET2 and TET3 synergistically regulate *Drosha* expression, by generating 5hmC across the gene body and by impacting chromatin accessibility. As DROSHA is involved in microRNA biogenesis, we proceed to investigate the impact of TET2/3 loss on microRNAs in iNKT cells. We report that among the downregulated microRNAs are members of the Let-7 family that downregulate *in vivo* the expression of the iNKT cell lineage specifying factor PLZF. Our data link TET proteins with microRNA expression and reveal an additional layer of TET mediated regulation of gene expression.

## Introduction

Ten Eleven Translocation (TET) proteins are enzymes that regulate the process of DNA demethylation by oxidizing 5 methylcytosine (5mC) to generate 5 hydroxymethylcytosine (5hmC) also known as the sixth base of our genome (1). In addition, TET proteins can oxidize 5hmC to generate additional modified cytosines, namely 5 formylcytosine (5fC) and 5 carboxylcytosine (5caC) (2, 3). The TET family of proteins consists of three members: TET1, that is most highly expressed in embryonic stem cells (ESCs), TET2, which is broadly expressed in various cell types and developmental stages, and TET3 that is more highly expressed as cells differentiate (4). All three TET proteins exert critical roles in shaping the development and function of a vast array of cells (5, 6). We have previously demonstrated that 5hmC is dynamically distributed across the genome of thymic T cell subsets (7). During the process of T cell lineage specification, 5hmC was shown to be increased in the gene body of very highly expressed genes and in active enhancers (7). Similar findings have been reported for murine and human peripheral T cells (7-10), indicating the critical role of TET proteins and 5hmC in regulating gene expression in T cells (5).

To dissect the *in vivo* roles of TET proteins in T cell development we generated *Tet2*-/-*Tet3*flx/flx CD4 cre (*Tet2/3* DKO) mice (11). We focused our analysis on concomitant deletion of TET2 and TET3 since our data indicated redundancy between TET proteins (11). The phenotype of the *Tet2/3* DKO mice was complex, revealing that TET proteins are critical regulators of various T cell types. Specifically, TET2 and TET3 are fundamental for the stability of the transcription factor (TF) FOXP3 and thus the functionality and stability of regulatory T cells (Tregs) (12). In addition, *Tet2/3* DKO mice exhibit a striking expansion of invariant natural killer (iNKT) T cells (11).

iNKT cells are an unconventional type of T cells that express an invariant TCR Vα14 chain and recognize lipids instead of peptides (13). iNKT cells develop in the thymus endowed already with effector properties and they have the ability to generate significant amount of cytokines, immediately upon antigen encounter (14). iNKT cell lineage commitment is orchestrated by the TF Promyelocytic leukaemia zinc finger (PLZF) protein, which endows iNKT cells with effector properties (15, 16). In the thymus, we can discern 3 subsets based on the expression of TFs and their functional properties: NKT2 express GATA3, NKT17 express RORγt and NKT1 express T-bet (17-20). iNKT cells exert important roles in recognition of bacterial pathogens and have been shown to be of clinical value in the context of cancer immunotherapy (14, 21-24). Thus, deciphering the molecular mechanisms that shape their differentiation and functionality is of outmost importance in order to take full advantage of their effector properties (25, 26).

We have previously demonstrated that *Tet2/3* DKO iNKT cells show increased expression of the TF RORγt (11, 27, 28). Integration of our genome wide datasets evaluating gene expression, whole genome methylation, whole genome hydroxymethylation and chromatin accessibility analysis in control and *Tet2/3* DKO iNKT cells revealed that TET2 and TET3, by regulating DNA demethylation, upregulate lineage specifying TFs such as T-bet and Th-POK that are critical for iNKT cell lineage diversification and for suppression of RORγt (11), in a TET2 dependent catalytic manner (29). However, not all the observed differences in the gene expression program of *Tet2/3* DKO iNKT cells can be attributed to gain of methylation in promoters or enhancers of the differentially expressed genes (11). That was particular true in the context of genes that were gaining expression upon loss of TET proteins (11). One possibility is that deletion of TET proteins can result in downregulation of repressors, allowing the upregulation of the targeted genes (30, 31). Repression of genes can occur by small RNAs that target mRNAs and can mediate their degradation (32). DROSHA regulates the generation of precursor miRNAs in the nucleus and then further processing occurs in the cytoplasm by DICER and the ARGONAUTE complex (33, 34). Notably, miRNAs are important for iNKT cell development as indicated by Dicer deficient mice (35, 36). In this study, we report that TET proteins regulate expression of *Drosha* in iNKT cells. We demonstrate that *Tet2/3* DKO iNKT cells show altered expression of precursor and mature miRNAs. Among the identified downregulated miRNAs are members of the Let-7 family that has been demonstrated *in vivo* to target and downregulate the transcription factor PLZF in iNKT cells (37).

## Results and Discussion

Analysis of our previously published RNA-seq datasets (11) revealed that *Drosha* was downregulated in *Tet2/3* DKO iNKT cells (**Figure 1A**). To further dissect the molecular mechanisms by which TET2 and TET3 can impact expression of *Drosha* we assessed 5hmC distribution across the gene body. 5hmC upon treatment with bisulfite sequencing is converted to cytosine-5-methylenesulfonate (CMS) (38). Analysis of CMS immunoprecipitation with sequencing (CMS-IP seq) (39, 40) datasets (11) revealed that in wild type iNKT cells 5hmC is distributed across the gene body of *Drosha* (**Figure 1B**). We have previously demonstrated that 5hmC is enriched in the gene body of highly expressed genes, whereas the promoters of these genes are devoid of 5hmC, in conventional T cells and unconventional iNKT cells (7, 11). Similar findings have been reported for naïve and helper T cell subsets (8-10) as well as for regulatory T cells (41). In addition, we have previously shown that 5hmC correlates with chromatin accessibility in both conventional and unconventional T cells (11, 29). We then investigated how loss of TET proteins may impact chromatin accessibility in the *Drosha* locus. Thus, we compared our datasets (11) for assay for transposase accessible chromatin with sequencing (ATAC-seq) (42) for wild type and *Tet2/3* DKO iNKT cells. We demonstrate that in *Tet2/3* DKO iNKT cells there is reduced accessibility in an intragenic genomic region (mm10: chr15:12,894,551-12,896,829) that has increased accessibility and enrichment of 5hmC in wild type iNKT cells (**Figure 1B**). Due to the low abundance of 5hmC in *Tet2/3* DKO thymic T cell subsets we were not able to perform CMS-IP seq for the *Tet2/3* DKO iNKT cells (11). However, we performed whole genome bisulfite sequencing (WGBS) in order to assess at single-nucleotide resolution the modification status of cytosine. Our analysis revealed a gain of methylation at this intragenic region in the *Drosha* locus at the *Tet2/3* DKO iNKT cells (**Figure 1B**).

**Figure 1.**
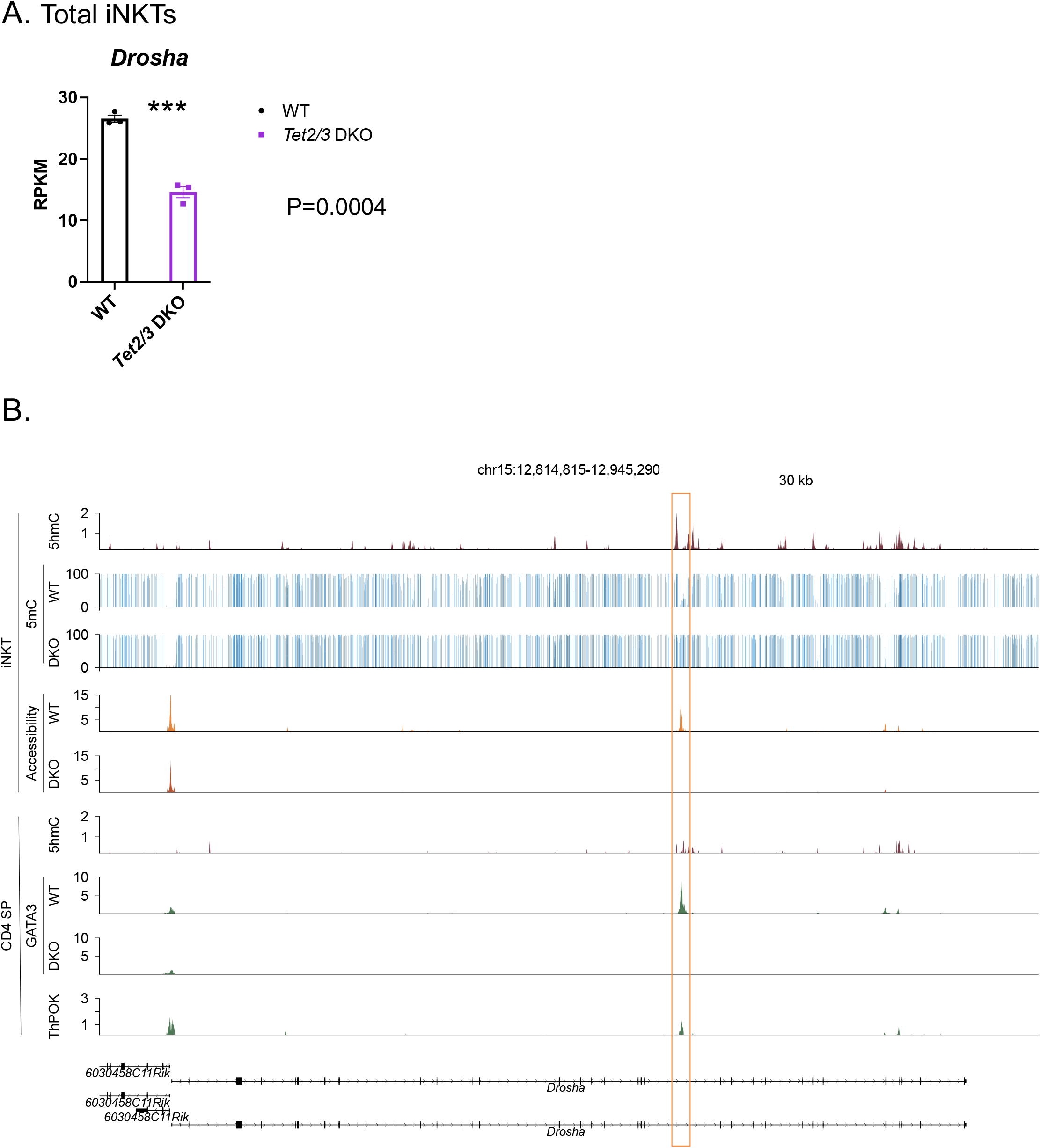
TET2 and TET3 regulate expression of *Drosha* in thymic iNKT cells. **A**. Gene expression of *Drosha* in WT (*in black*) and *Tet2/3* DKO thymic iNKT cells (*in purple*), evaluated by RNA-seq. 3 biological replicates per genotype were assessed. *** (p =0.0004), unpaired t test. Each dot represents an individual biological replicate. Horizontal lines indicate the mean (s.e.m.). **B**. Portraits of epigenetic regulation (determined by 5hmC, 5mC and chromatin accessibility) in thymic iNKT cells and transcriptional regulation in CD4 SP cells of the *Drosha* locus. Genome browser view of 5hmC distribution (by CMS-IP seq) in the gene body of *Drosha* in WT iNKT cells reveals enrichment of this modification indicating TET activity. 2 biological replicates were analyzed. Evaluation of 5mC by WGBS in WT and *Tet2/3* DKO iNKT cells. Note the gain of methylation in *Tet2/3* DKO iNKT cells in the indicated region that coincides with enrichment of 5hmC at WT iNKT cells (mm10: chr15:12,894,551-12,896,829). 2 biological replicates of WGBS per genotype were evaluated. Portraits of chromatin accessibility (assessed by ATAC-seq) of the *Drosha* locus in WT and *Tet2/3* DKO thymic iNKT cells. Peaks indicate accessibility. Note the loss of accessibility in the *Tet2/3* DKO iNKT cells at the indicated site that is enriched for 5hmC in the WT iNKT cells. 3 biological replicates per genotype were evaluated. 5hmC distribution (determined by CMS-IP seq) in the gene body of *Drosha* in WT CD4 SP cells reveals enrichment of this modification in the same intragenic site. 2 biological replicates were evaluated. GATA3 binds in this potentially regulatory site in WT CD4 SP cells as determined by CUT&RUN, whereas GATA3 binding is not detected in *Tet2/3* DKO CD4 SP cells. 3 biological replicates for WT CD4 cells and 2 biological replicates for *Tet2/3* DKO were analyzed. ThPOK binding is detected by ChIP-seq in WT CD4 SP cells at the indicated intragenic site. 3 biological replicates were evaluated. The arrows indicate the direction of transcription.

We hypothesize that this intragenic region may exert regulatory function to promote the expression of *Drosha*. We have previously shown that 5hmC decorates active enhancers (7). Additional studies have demonstrated a strong correlation of 5hmC with active enhancers in various T cell subsets (8, 41). In many cases these regulatory elements that require 5hmC enrichment in order to be active are intragenic, such as the CNS2 enhancer in the *Foxp3* locus (12, 43, 44), an intragenic enhancer that regulates stable expression of *Cd4* gene in CD4 cells (45) as well as the proximal enhancer of *Zbtb7b* gene that encodes Th-POK (29). We have also demonstrated that 5hmC decorates intragenic cite A at the *Zbtb7b* locus to regulate the accessibility and the binding of the transcription factor GATA3 (29). It has been previously suggested that the binding of GATA3 to cite A promotes Th-POK expression (46). We have previously discovered a shared gene expression program between *Tet2/3*

DKO thymic iNKT cells and CD4 single positive (SP) cells (29). As DROSHA is expressed in both subsets we investigated if its expression was also affected in CD4 SP cells. We report that *Drosha* is downregulated in *Tet2/3* DKO CD4 SP cells (**Supplementary Figure 1**). Interestingly, there is 5hmC enrichment at the same intragenic site of the locus in WT CD4 SP cells (**Figure 1B**). In addition, we looked into our data assessing recruitment of GATA3 (by CUT&RUN) in WT and *Tet2/3* DKO CD4 SP cells (29). We discover that GATA3 binds in this region in WT CD4 SP cells, whereas no binding was detected in *Tet2/3* DKO CD4 SP cells. Moreover, we looked into the binding of Th-POK by using publicly available ChIP-seq datasets (47) and we demonstrate binding of Th-POK in this potentially regulatory region in CD4 SP cells. Collectively, our findings suggest that TET2 and TET3 generate 5hmC and regulate chromatin accessibility in the *Drosha* locus to promote the expression of the gene (**Figure 1**). Further studies are required to elucidate the precise regulatory elements that control the expression of *Drosha*. However, as TET2 and TET3 deletion results in partial reduction of *Drosha* expression and not complete loss it becomes apparent that additional mechanisms are in place to control the expression of this gene.

As DROSHA is involved in regulating the pathway of microRNAs (miRNAs) we asked whether the reduced expression of *Drosha* has an impact on the miRNAs that are expressed in *Tet2/3* DKO iNKT cells. To identify small RNAs that are impacted we isolated thymic iNKT cells by FACS sorting (**Supplementary Figure 2**) from wild type or *Tet2/3* DKO mice and we performed small RNA-seq (**Figure 2A**). Comparison of precursor and mature miRNAs in the WT and the *Tet2/3* DKO iNKT samples confirmed that samples of the same genotype were similar to each other (**Supplementary Figure 3**). Our analysis compared expression of precursor (**Supplementary Table 1**) and mature miRNAs (**Supplementary Table 2**) and we found that among those that were differentially expressed the majority were downregulated (**Figure 2B, C**). This could be due to the downregulation of *Drosha* expression. We then focused on the affected mature miRNAs (**Figure 2C**). The vast majority of the differentially expressed mature miRNAs were downregulated in the *Tet2/3* DKO iNKT cells (**Figure 2C**). An additional mechanism could be that in the absence of TET proteins at least some miRNAs could gain cytosine methylation, resulting in their downregulated expression. However, when we looked into our previously generated WGBS data (11) we did not notice significant changes in methylation for the vast majority of the miRNAs that were differentially expressed in thymic iNKT cells. We only detected some gain of methylation in *mir199b* and *mir7058* (**Supplementary Figure 4**). Our analysis demonstrated that among the downregulated miRNAs were members of the Let-7 family. Specifically, we observed downregulation of Let-7c, Let-7b and Let-7k (**Figure 2C**). Interestingly, Let-7 miRNAs have been previously shown to target *Zbtb16* mRNA, which encodes for PLZF, for degradation in murine iNKT cells *in vivo* (37). Thus, we hypothesize that the downregulation of some of the members of the Let-7 family could result in increased expression of PLZF. We evaluated PLZF levels in WT and *Tet2/3* DKO iNKT cells by Flow cytometry (**Figure 3A**). Our data indicates that *Tet2/3* DKO iNKT cells exhibit upregulation of PLZF (**Figure 3A, B**). Thus, we propose that in *Tet2/3* DKO iNKT cells downregulation of some of the Let-7 miRNAs results in reduced targeting for degradation of *Zbtb16* mRNA, resulting in increased expression of PLZF (**Figure 3C**).

**Figure 2.**
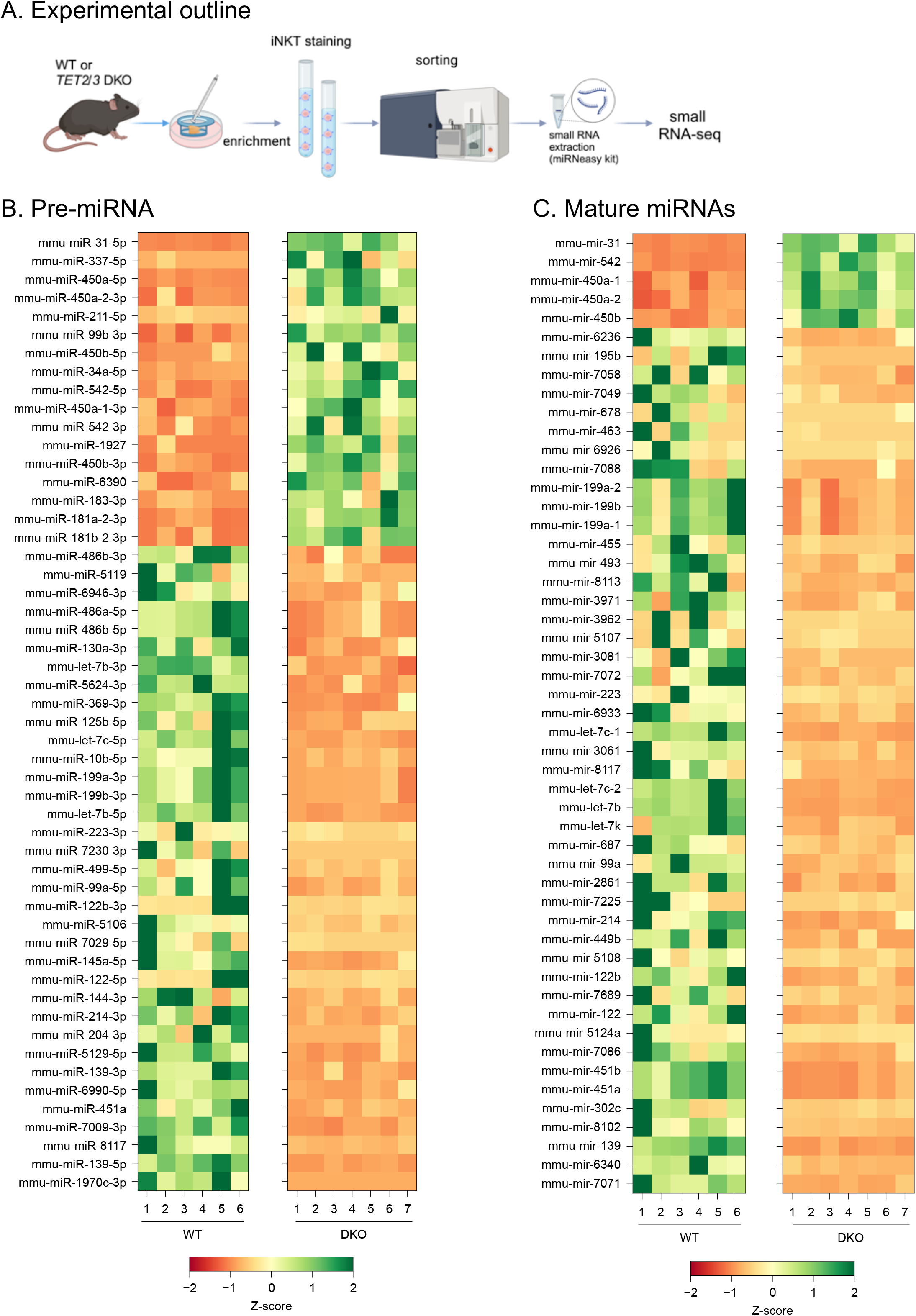
Differential expression of precursor (hairpin) and mature miRNAs in WT and *Tet2/3* DKO thymic iNKT cells. **A**. Experimental outline. **B**. Heatmap indicating hairpin miRNAs whose adjusted p-value < 0.05 and absolute log2 fold-change > 2. The z-score normalized expression values are shown. **C**. Heatmap indicating mature miRNAs whose adjusted p-value < 0.05 and absolute log2 fold-change > 2. The z-score normalized expression values are shown. Both male and female mice were used for each genotype. N=6 WT mice and N=7 *Tet2/3* DKO mice were used.

**Figure 3.**
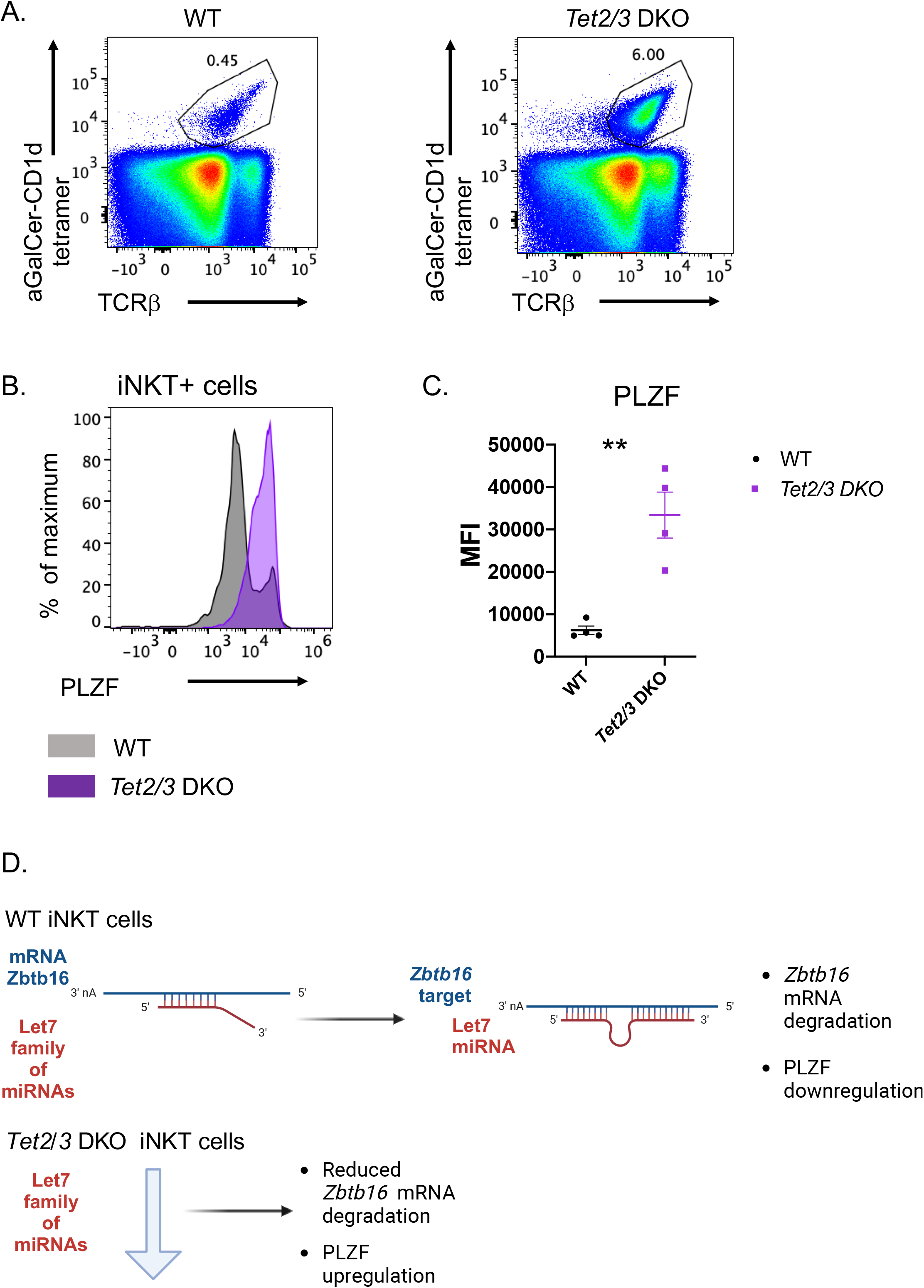
Let-7 miRNAs downregulation in *Tet2/3* DKO thymic iNKT cells contributes in upregulation of PLZF. **A**. Representative flow cytometry plots of thymocytes isolated from wild type and *Tet2/3* DKO mice identify iNKT cells as aGalCer-loaded tetramer^+^ and TCRβ intermediate cells. Representative histogram for the lineage specifying transcription factor PLZF indicates increased expression, determined by intracellular staining and Flow cytometry, in the *Tet2/3* DKO thymic iNKT samples (*in purple*) compared to WT (*in black*) counterparts. **B**. Plot comparing the median fluorescence intensity (MFI) of PLZF expression in WT (*in black*) and *Tet2/3* DKO (*in purple*) iNKT cells. 4 biological replicates per genotype were assessed. ** (p =0.0004), unpaired t test. Each dot represents an individual biological replicate. Horizontal lines indicate the mean (s.e.m.). **C**. Model for TET mediated regulation of members of the Let-7 miRNA family to impact PLZF expression in thymic iNKT cells.

## Conclusions

In this study, we report that TET2 and TET3 regulate the expression of *Drosha*. We also discover various miRNAs that are differentially expressed including downregulation of Let-7 miRNAs. However, we must emphasize that the NKT17 skewing of the *Tet2/3* DKO iNKT cells can be fully rescued by deletion of ThPOK and partially rescued by deletion of T-bet as we have previously shown (11). Importantly, our unbiased, integrative analysis of genome wide datasets indicated that both ThPOK and Tbet are targets of TET proteins based on 5hmC enrichment and gain of methylation upon concomitant TET2 and TET3 loss (11). Thus, in support of our previous findings that TET proteins exert multifaceted roles in regulating gene expression (30, 31, 48), we propose an additional layer of TET-mediated regulation of lineage specification by affecting expression of miRNAs.

## Methods

### Mice

Mice were housed in pathogen free conditions in the Genetic Medicine Building at University of North Carolina (UNC) Chapel Hill in a facility managed by the Division of Comparative Medicine at UNC Chapel Hill. All the experiments using mice in this study were performed according to our approved protocol by the UNC Institutional Animal Care and Use Committee (protocol no: 22-252). Age and sex-matched mice were analyzed. Male and female mice were used for our experiments. Control (C57BL/6 (B6), strain number: 000664), RRID: IMSR_JAX: 000664) mice were purchased from Jackson (Jax) laboratories and were bred in our facility at UNC. Tet2-/-Tet3flx/flx CD4 cre mice have been previously described (11, 29). Briefly, Tet2-/-mice (49) (Jax strain no; 023359, RRID: IMSR_JAX:023359) were crossed with Tet3flx/flx (50, 51) (Jax strain no: 031015, RRID: IMSR_JAX:031015) CD4cre mice (52). To determine the genotype of the mice, tissue was isolated and genomic DNA was extracted using Phire Animal Tissue Direct PCR kit (Thermo scientific, cat no F-140WH) following the manufacturer’s protocol. Then DNA fragments were amplified by PCR using the Phire DNA polymerase (Thermo scientific, cat no F-140WH) and specific primers using Biorad T100 or Biorad C1000 Touch thermocyclers.

### Cell preparation

Thymocytes were isolated from young mice 21-25 days old. Thymocytes were dissociated to prepare single cell suspensions as previously described (53, 54).

### Flow cytometry

Thymocytes were stained with PBS-57 loaded tetramer PE (dilution 1:400, from NIH tetramer core), TCRβ-PERCP/Cy5.5 (dilution 1:200, Biolegend, clone: H57-597, RRID: AB_1575173) and dead cells were excluded by using a live/dead dye efluor 780 (dilution 1:1000, eBioscience, cat. #65-0865-18) in FACS buffer (2% FBS in PBS) as described (54). Intracellular staining for PLZF AlexaFluor 647(dilution 1:100, BD Pharmingen, clone: R17-809, RRID: AB_2738238) was performed using the Foxp3 Transcription factor staining buffer set (eBioscience, cat. no:00-5523-00)(54). Samples were analyzed in a Novocyte 3005 Flow Cytometer (Agilent) using NovoExpress software (Agilent). Subsequently, the acquired data were analysed and plots were generated using FlowJo (Treestar).

### FACS sorting

Total thymocytes were stained with biotinylated mouse anti-CD24 (Biolegend, clone:M1/69, RRID: AB_312837). CD24+ cells were depleted using mouse streptavidin magnetic beads (anti-mouse Rapidspheres, cat no 19860, STEMCELL Technologies) following the manufacturer’s guidelines as previously described (29, 53). Enriched cells were stained with efluor 780 viability dye (dilution 1:1000, eBioscience, cat. #65-0865-18), aGalactosyl-Ceramide loaded tetramer (conjugated with PE, obtained from NIH tetramer core, dilution 1:400), TCRβ-PERCP/Cy5.5 (dilution 1:200, Biolegend, clone: H57-597, RRID: AB_1575173). Live TCRβ^+^, tetramer^+^ cells were sorted and used to isolate RNA. The purity of the samples after sorting was >98%. The cells were sorted using either a FACSAria II or a FACSymphony S6 Sorter (Becton Dickinson).

### Statistical Analysis

For the statistical analysis we used Prism software (Graphpad). We applied unpaired student’s *t* test. In the relevant figure legends, we indicated p-values for statistically significant differences (p < 0.05). Data are mean ± s.e.m. In the graphs, each dot represents a mouse. Unless otherwise indicated the p-value was not statistically significant. Differences were considered significant when p < 0.001 (^∗∗∗^); < 0.0001 (^∗∗∗∗^). Both male and female mice from different litters were evaluated, with reproducible results.

### RNA isolation, Library preparation of small RNAs and sequencing

FACS sorted iNKT cells were lysed in RLT plus lysis buffer from the miRNeasy plus kit (Qiagen, cat no: 217084). Total RNA was isolated following the instructions provided by the manufacturer and was quantified using Qubit RNA High Sensitivity assay (Invitrogen) in Qubit 4 Fluorometer (Invitrogen). Total RNA was provided to the UNC High Throughput Sequencing Facility (HTSF). RNA integrity was evaluated with a Tapestation (Agilent) using High Sensitivity RNA ScreenTape (Agilent). RNA with RIN value>9 was used for library preparation. Small RNA libraries were generated using the Revvity NETFLEX small RNA sequencing kit V4. Libraries were pooled and sequenced in an Illumina NextSeq2000 P1 Single End 1×50 to obtain 100 million single end reads. 6 biological replicates for wild type and 7 biological replicates for DKO samples were analyzed. Both male and female mice were evaluated.

### Datasets used in this study

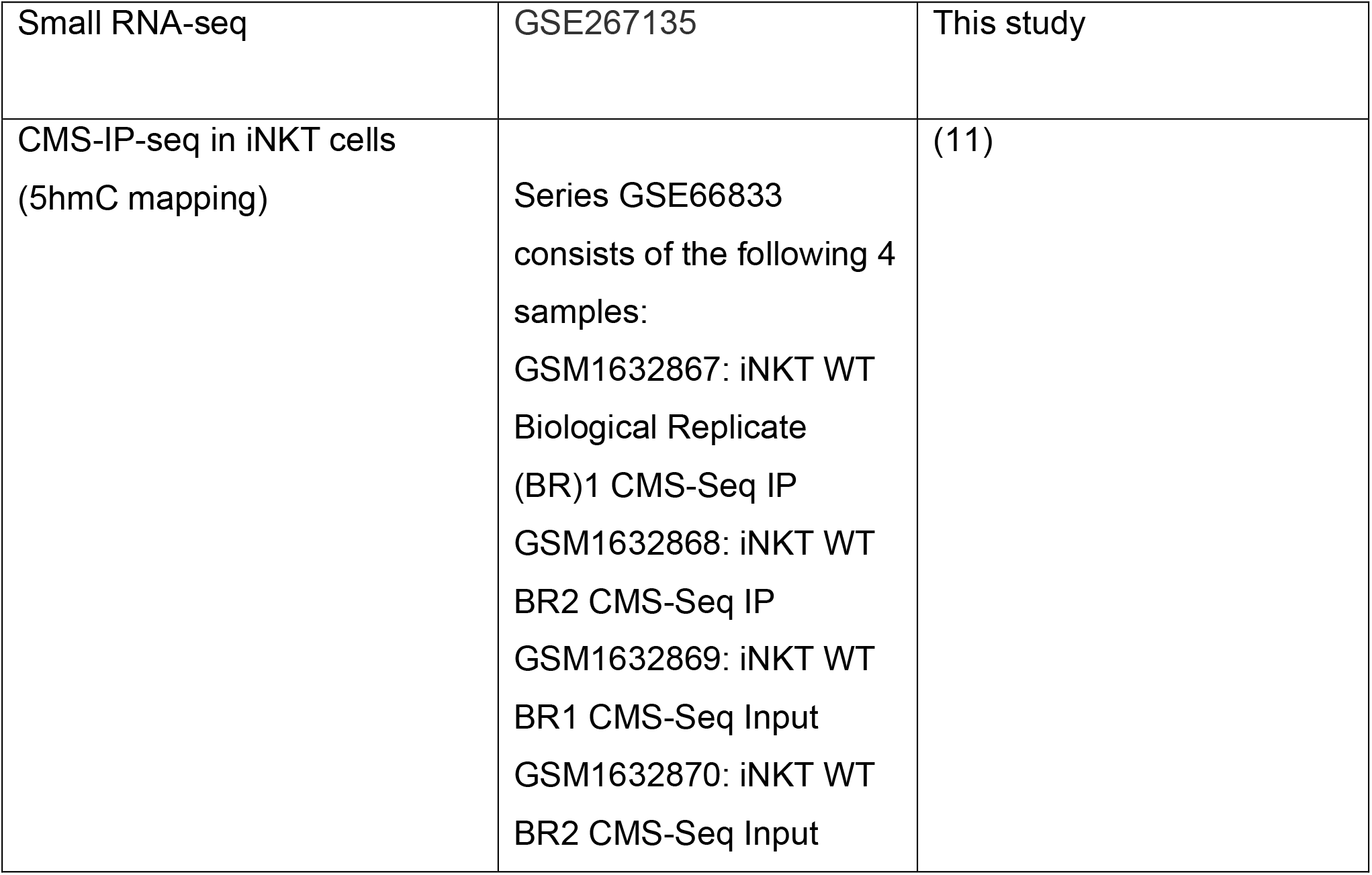
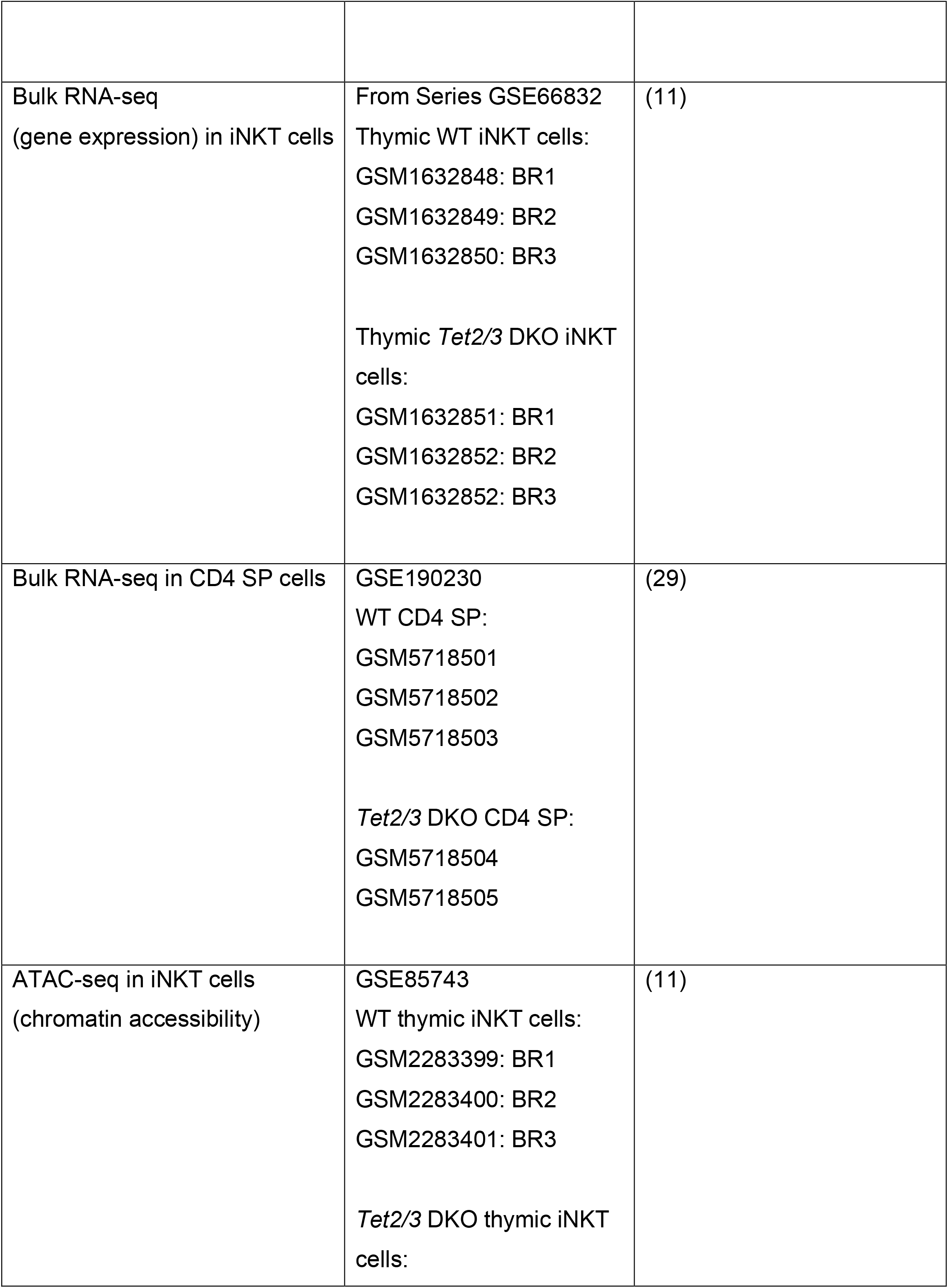
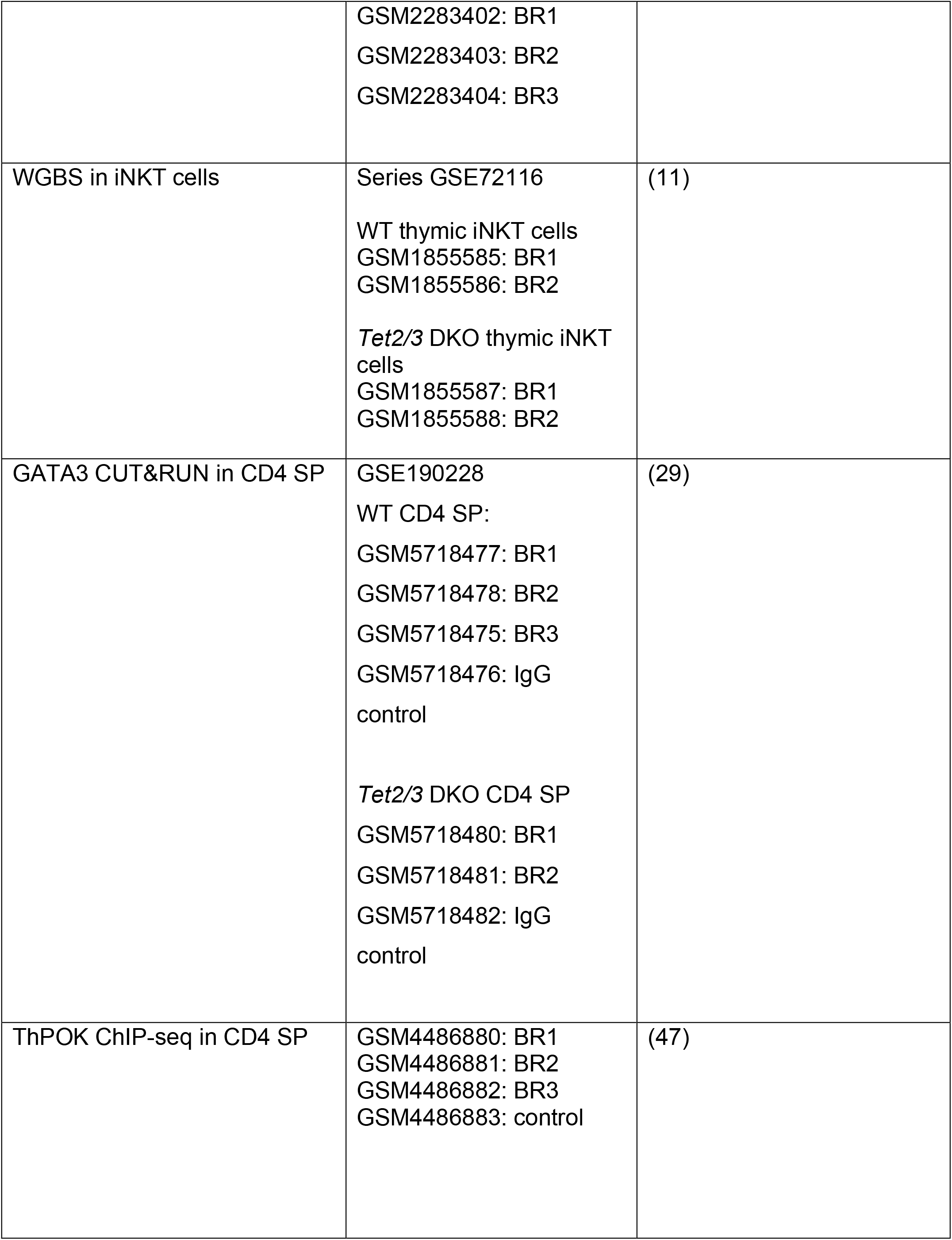

#### Small RNA-seq data analysis

The small RNA samples were processed using nf-core/smrnaseq (2.2.4) using default parameters (55, 56). The differential expression analysis was done using nf-core/differentialabundance (1.4.0) using default parameters (57).

#### CMS-seq data analysis

The CMS-IP and input reads from 2 biological replicates of WT iNKT cells were mapped against mm10 using Bismark (0.22.3) (58). The mapping was done using the Bowtie 2 (2.4.1) (59) backend in the paired-end mode with the following parameter values: -I 0 -X 600 -N 0. The coverage tracks were generated using HOMER (4.10) (makeBigWig.pl -norm 1e6) (60).

#### ATAC-seq data analysis

Adapter trimming and quality filtering of the sequencing libraries (3 biological replicates per genotype, 6 samples in total) was done using fastp (0.21.0)(61) with the default parameters. The sequencing libraries were mapped against mm10 using Bowtie 2 (2.4.1) (–very-sensitive -X 2000)(59). Mitochondrial reads were removed after alignment. Additional filtering was done using samtools (1.12)(62) using the following parameter values: -q 30 -h -b -F 1804 -f 2. Reads with identical sequences were filtered and only one was retained for subsequent analysis. The coverage tracks were generated from the samples obtained by pooling the biological replicates using HOMER (4.10) (makeBigWig.pl -norm 1e6)(60).

WGBS, CUT&RUN and ChIP-seq data analysis has been previously described (29).

## Figure Legends

**Supplementary Figure 1: TET2 and TET3 regulate expression of *Drosha* in thymic CD4 SP cells. A**. Gene expression of *Drosha* in WT (*in black*) and *Tet2/3* DKO thymic iNKT cells (*in purple*), evaluated by RNA-seq. 3 biological replicates for WT and 2 biological replicates for Tet2/3 DKO CD4 SP cells were assessed. *** (p =0.0003), unpaired t test. Each dot represents an individual biological replicate. Horizontal lines indicate the mean (s.e.m.).

**Supplementary Figure 2**: **Sorting strategy and purity assessment of FACS sorted iNKT cells. A**. Representative flow cytometry plots indicating the gating selection to exclude doublets and isolate live (LD APC/CY7 negative), wild type iNKT cells (aGalCer loaded tetramer positive, TCRβ intermediate cells) by FACS sorting. **B**. FACS plots indicating purity of a representative sample of wild type iNKT cells after FACS sorting. **C**. As in **A**. for *Tet2/3* DKO iNKT sample. **D**. FACS plots indicating purity of a representative sample of *Tet2/3* DKO iNKT cells after FACS sorting.

**Supplementary Figure 3**: **Evaluating similarity of RNA samples. A**. Dendrograms indicating the clustering of precursor miRNAs and **B**. mature miRNAs samples.

**Supplementary Figure 4**: **Methylation portraits in mature miRNAs**. Assessing cytosine methylation by WGBS revealed some gain of methylation in *Tet2/3* DKO iNKT cells in two of the mature miRNAs that were downregulated in *Tet2/3* DKO iNKT cells. 5mC distribution in WT and *Tet2/3* DKO iNKT cells for A. *Mir199b* and B. *Mir7058*.

**Supplementary Table 1**: Results of differential gene expression analysis of hairpin miRNAs in wild type and *Tet2/3* DKO iNKT cells.

**Supplementary Table 2**: Results of differential gene expression analysis of mature miRNAs in wild type and *Tet2/3* DKO iNKT cells.

## Data availability statement

The datasets generated for this study have been deposited in the Gene Expression Omnibus (GEO) public repository with the accession code: GSE267135.

## Author Contributions

MG performed cell isolation, Flow cytometry, prepared samples for FACS sorting, isolated RNA and contributed in maintaining relevant mouse colonies. TÄ analyzed RNA-seq, CMS-IP seq, WGBS, CUT&RUN, ChIP-seq and small RNA-seq data, generated figures and wrote the relevant method sections. JEV and JBB assisted in Flow cytometry experiments and mouse colony management. AT conceived the study, performed experiments, analyzed data, supervised research, acquired funding and wrote the manuscript. All authors agree with the content of the manuscript.

## Acknowledgments

We acknowledge Ms. Theresa Hegarty (UNC DCM Colony Management Core) for excellent mouse colony management. We thank Ms. Janet Dow, Mr. Roman Bandy and Ms. Ayrianna Woody of the UNC Flow Cytometry Core for FACS sorting. We thank the UNC HTSF for preparing small RNA-seq libraries and for performing sequencing. We are grateful to the NIH tetramer core for generously providing PBS-57 loaded mouse CD1d tetramers. Some of the images were generated with Biorender.

