## Supplementary figures for "TET proteins regulate Drosha expression and impact microRNAs in iNKT cells"

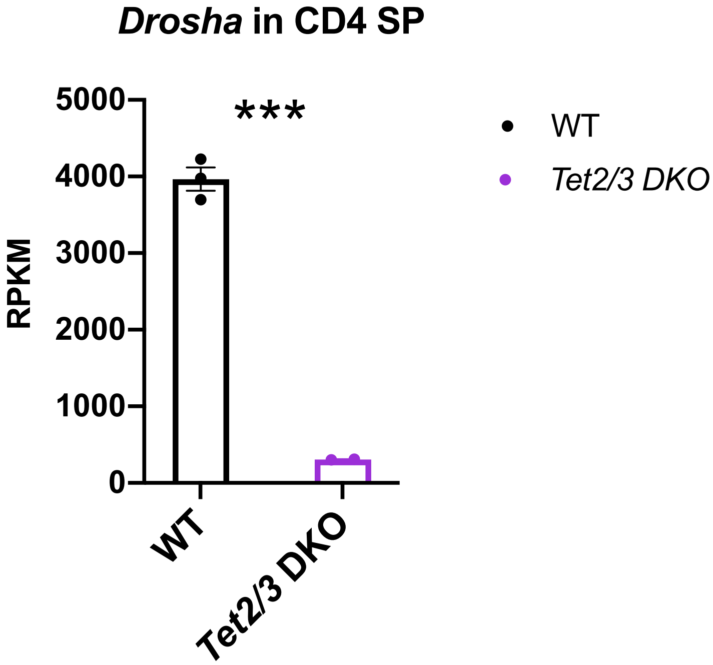


Supplementary Figure 1

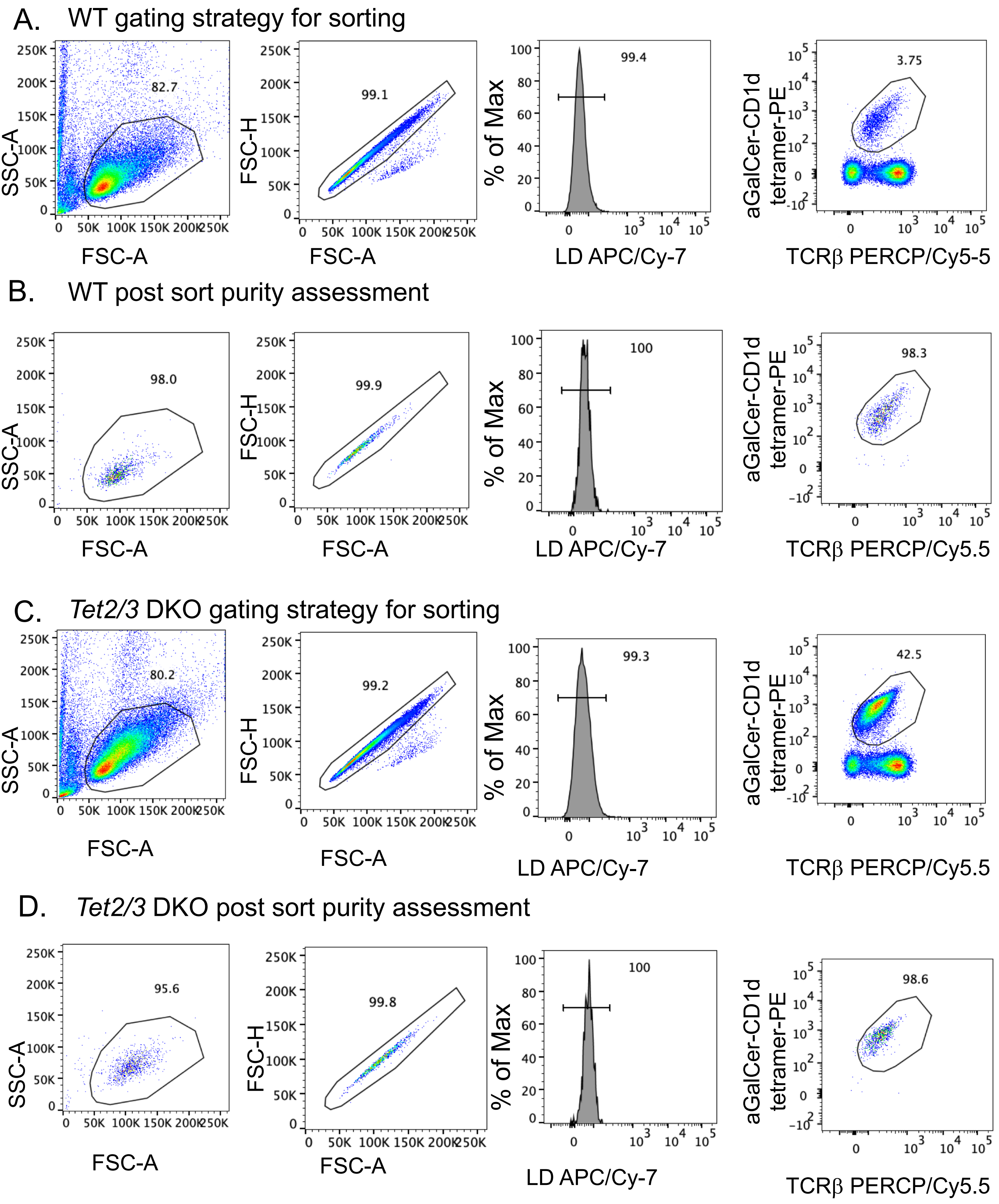


Supplementary Figure 2

A. Sample clustering dendrogram, 500 most variable genes

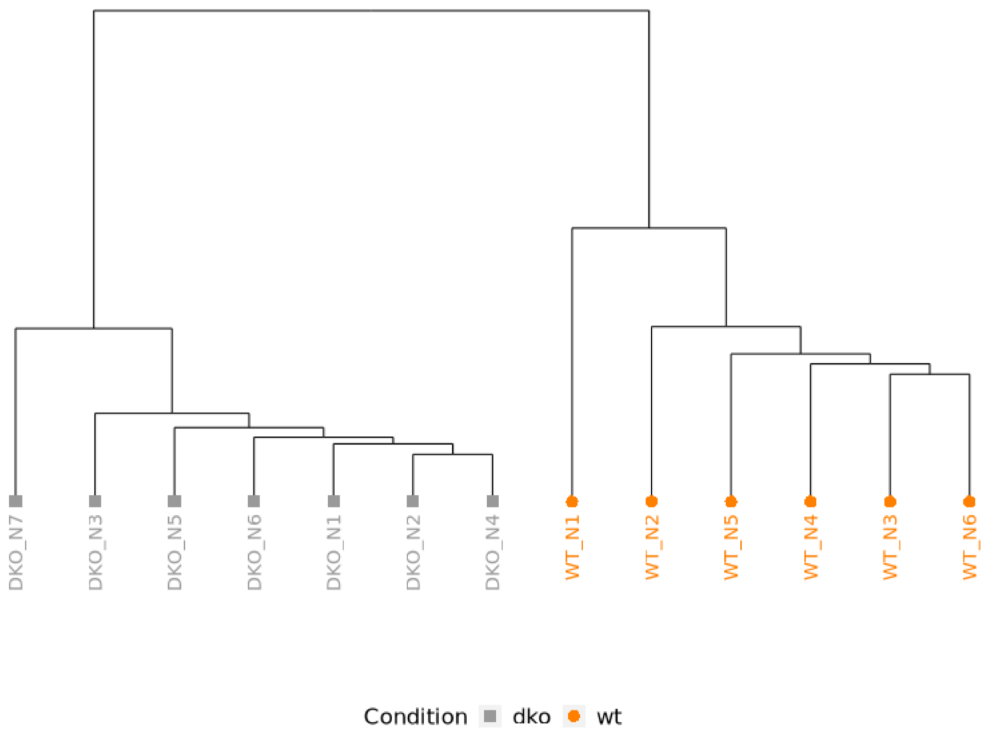


B. Sample clustering dendrogram, 500 most variable genes

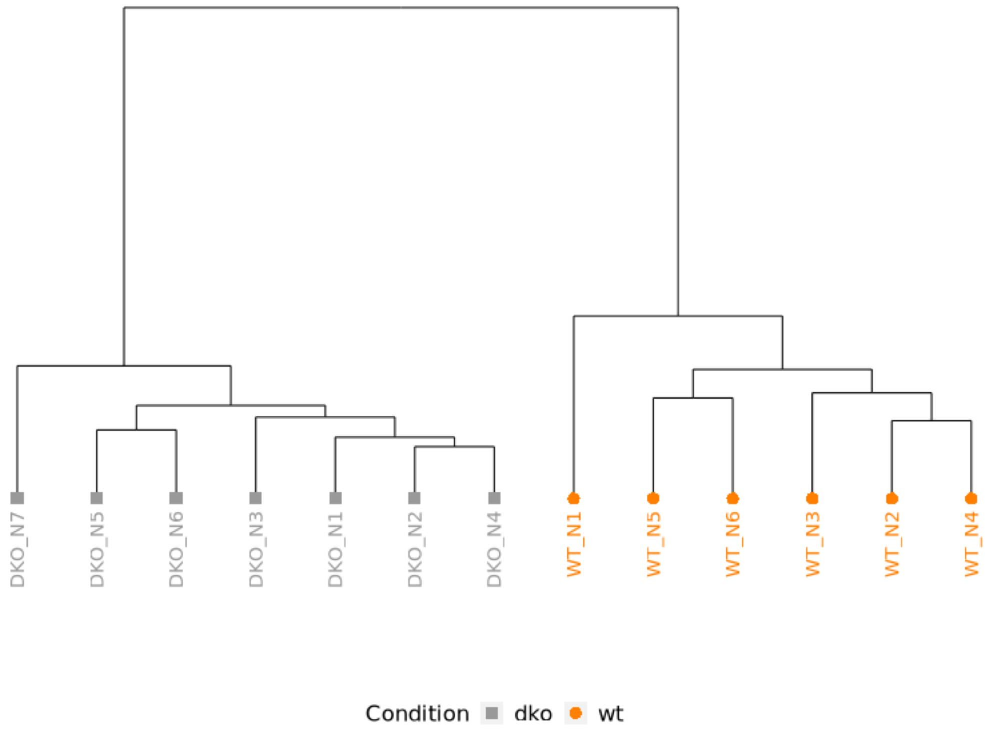


Supplementary Figure 3

A. Mir199b

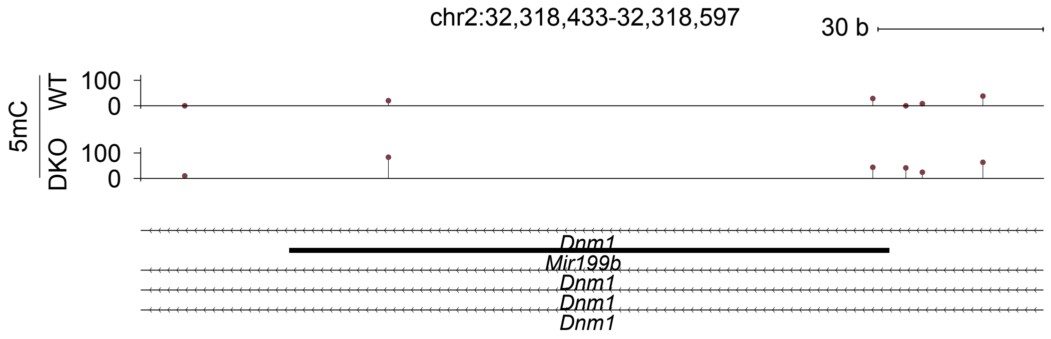


B. Mir7058

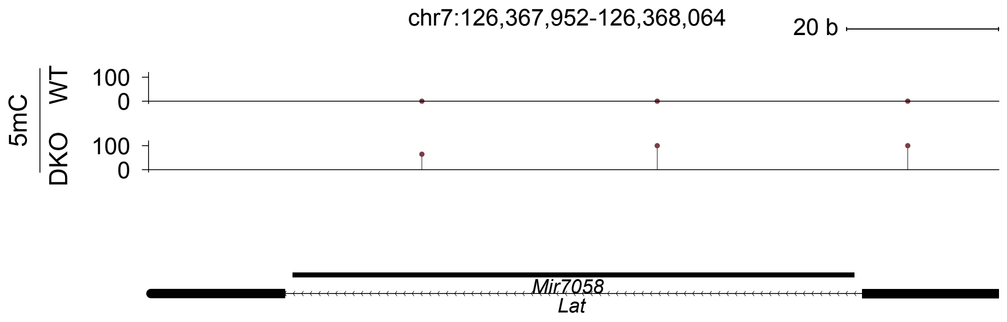
